# Pathogen and human NDPK-proteins promote AML cell survival via monocyte NLRP3-inflammasome activation

**DOI:** 10.1101/2023.01.02.522534

**Authors:** Sandro Trova, Fei Lin, Santosh Lomada, Matthew Fenton, Bhavini Chauhan, Alexandra Adams, Avani Puri, Alessandro Di Maio, Thomas Wieland, Daniel Sewell, Kirstin Dick, Daniel Wiseman, Deepti P Wilks, Margaret Goodall, Mark T Drayson, Farhat L Khanim, Christopher M Bunce

**Affiliations:** School of Biosciences, University of Birmingham, UK; Institute of Experimental and Clinical Pharmacology and Toxicology, Heidelberg University, Mannheim, Germany; Division of Cancer Sciences. University of Manchester, UK; Manchester Cancer Research Centre Biobank, Cancer Research UK Manchester Institute, The University of Manchester, UK; Institute of Immunology and Immunotherapy, University of Birmingham, UK; Clinical Sciences, University of Birmingham, UK

**Author notes:** Corresponding Author: Chris Bunce, School of Biosciences, University of Birmingham, Edgbaston, Birmingham, UK. B15 2TT. Equal senior authors (Farhat L Khanim, Christopher M Bunce). Deceased (Margaret Goodall).

## Abstract

A history of infection has been linked with increased risk of acute myeloid leukaemia (AML) and related myelodysplastic syndromes (MDS). Furthermore, AML and MDS patients suffer frequent infections because of disease-related impaired immunity. However, the role of infections in the development and progression of AML and MDS remains poorly understood. We and others previously demonstrated that the human nucleoside diphosphate kinase (NDPK) NM23-H1 protein promotes AML blast cell survival by inducing secretion of IL-1β from accessory cells. NDPKs are an evolutionary highly conserved protein family and pathogenic bacteria secrete NDPKs that regulate virulence and host-pathogen interactions. Here, we demonstrate the presence of IgM antibodies against a broad range of pathogen NDPKs and more selective IgG antibody activity against pathogen NDPKs in the blood of AML patients and normal donors, demonstrating that *in vivo* exposure to NDPKs likely occurs. We also show that pathogen derived NDPK-proteins faithfully mimic the catalytically independent pro-survival activity of NM23-H1 against primary AML cells. Flow cytometry identified that pathogen and human NDPKs selectively bind to monocytes in peripheral blood. We therefore used vitamin D_3_ differentiated monocytes from wild type and genetically modified THP1 cells as a model to demonstrate that NDPK-mediated IL-1β secretion by monocytes is NLRP3-inflammasome and caspase 1 dependent, but independent of TLR4 signaling. Monocyte stimulation by NDPKs also resulted in activation of NF-κB and IRF pathways but did not include the formation of pyroptosomes or result in pyroptotic cell death which are pivotal features of canonical NLRP3 inflammasome activation. In the context of the growing importance of the NLRP3 inflammasome and IL-1β in AML and MDS, our findings now implicate pathogen NDPKs in the pathogenesis of these diseases.

**Author Summary:** Acute myeloid leukaemia (AML) and myelodysplastic syndromes MDS) are related blood cancers that are associated with frequent infections because the cancers suppress normal immunity. These infections are therefore generally considered as medical complications arising as a result of but separate to the cancer. However, we provide evidence here that infections may promote or drive cancer progression. We and others previously demonstrated that a human protein called NM23-H1 promotes the survival of AML cells by eliciting survival signals from other cells. NM23-H1 belongs to a highly conserved family of proteins that also occur in bacteria and fungi that cause infections in AML and MDS patients. Here we demonstrate that these bacterial and fungal proteins recapitulate the pro-survival effect of NM23-H1 on AML cells. We also determine that these effects are mediated via mechanisms already known to be important in the development and progression of AML and MDS. This study is the first to identify NM23-H1 like proteins from pathogenic microorganisms as novel activators of these pathways. These findings have important implications for how we understand infections in AML and MDS patients and suggest that in addition to being the consequence of these diseases, infections may also drive the cancer process.

## Introduction

Population based studies have identified that a history of infection associates with increased risk of both acute myeloid leukaemia (AML) and the related myelodysplastic syndromes (MDS) (1) which also have a high risk of progression to AML. Infection related risk remains significant even when confined to infections occurring three or more years before AML/MDS diagnosis (1). Neutropenic sepsis is also the most common route to AML presentation and a major cause of death in both AML and MDS (2-5). Following diagnosis, infections continue to represent key clinical challenges in AML/MDS treatment and management. It is generally accepted that increasing immune impairment arising as a consequence of disease progression is responsible for the frequency of life-threatening infections in these patients. However, the reciprocal role of infections in the development and progression of AML and MDS remains poorly understood and largely ignored.

Despite this, the important role of the inflammatory cytokine interleukin-1β (IL-1β) in AML has become increasingly recognized. Early studies demonstrated that soluble IL-1 receptor (sIL-1R) and IL-1β receptor antagonists (IL-1RA) reversed the pro-survival effect of IL-1β on AML blasts *in vitro* (6). More recently, a functional screen of 94 cytokines identified that IL-1β elicited expansion of myeloid progenitors whilst suppressing the growth of normal progenitors in 67% of AML patients (7). In the same study, silencing of the IL-1 receptor led to significant suppression of clonogenicity and *in vivo* disease progression (7). Similarly chronic IL-1β exposure triggers the selective *in vivo* expansion of *Cebpa*-deficient multipotent hematopoietic progenitors, demonstrating a role for inflammation in selecting early premalignant clones in the bone marrow (8). A recent study also identified that AML patients with low IL-1β along with high levels of the IL-1 receptor antagonist IL-1RA were protected against relapse following immunotherapy, further implicating inflammation in promoting relapse (9).

IL-1β secretion is regulated by activation of multiprotein complexes called inflammasomes. The *IL1B* gene encodes an inactive cytoplasmic pro-IL-1β protein that is cleaved upon inflammasome activation by caspase-1, to form the active secreted from of IL-1β. Several studies have implicated the NLRP3-inflammasome in AML progression(10-12). Equally, the importance of NLRP3 activation in the pathogenesis of myelodysplastic syndromes (MDS) is well established (13-16). The NLRP3 inflammasome (comprised of the NOD-, leucine-rich repeat (LRR)-and pyrin domain (PYD)-containing protein 3 (NLRP3), the adapter apoptosis-associated speck-like protein containing a caspase recruitment domain (ASC), and the effector protease caspase-1) orchestrates Toll-like receptor 4 (TLR4) responses to both sterile and infection-related inflammation. Sterile inflammation occurs in response to host derived damage associated molecular pattern (DAMP) signals (including S100A9 (S100 Calcium Binding Protein A9)) (13, 17), whereas infection-related inflammation is mediated by pathogen associated molecular pattern (PAMP) signals (classically LPS (lipopolysaccharide) from Gram-negative bacteria) (18, 19).

Following on from reports that elevated plasma levels of the human nucleoside diphosphate kinase (NDPK) NM23-H1 are associated with poor prognosis in AML (20-22), we and others demonstrated that exogenous NM23-H1 indirectly promotes AML blast cell survival via the induction of inflammatory cytokines including IL-β (23-25). NDPK proteins (also termed NDK, NME) are evolutionarily highly conserved across prokaryotes and eukaryotes. The most studied function of NDPKs is catalysis of the reversible γ-phosphate transfer from nucleoside triphosphates (NTPs) to NDPs (26). However, microbial NDPKs also have complex roles in virulence (27, 28), regulating immune responses to infection (28, 29) and host-pathogen interactions (30-32). Therefore, it is now important to consider whether, in addition to NM23-H1 driving AML progression, infection derived NDPKs may also directly exacerbate risk of AML and MDS and disease progression.

We demonstrate here that NDPKs from pathogens commonly associated with infections and sepsis in AML and MDS patients, faithfully recapitulate the activity of NM23-H1 in promoting primary AML cell survival. We further demonstrate that human and pathogen NDPKs induce IL-1β production in monocytes via an NLRP3 and caspase-1 dependent but TLR4-independent mechanism. Monocyte stimulation by NDPKs also resulted in activation of NF-κB and interferon regulatory factor (IRF) signaling pathways, but did not include the formation of pyroptosomes or pyroptotic cell death, which are pivotal features of canonical NLRP3 inflammasome activation (33, 34). Thus, our data indicate a novel role for pathogen derived NDPK proteins in promoting disease progression in AML and MDS.

## Results

### NM23-H1 and pathogen NDPKs are highly conserved

We aligned the human NM23-H1 amino acid sequence with publicly available amino acid sequence data for NDPKs from four bacterial strains and one fungal strain commonly associated with infections in AML and MDS patients, namely, *Escherichia coli (E.coli), Staphylococcus aureus (S.aureus), Streptococcus pneumoniae (S.pneumoniae), Klebsiella pneumoniae (K.pneumoniae), and Candida albicans C.albicans* (Figure 1A). Sequence conservation varied from 42-64 % (supplementary Table S1) and included conservation of key residues (red stars) required for enzymatic and oligomerization functions in NM23-H1 (Figure 1A). We also used available crystal structures to compare the tertiary structures of NM23-H1 with NDPKs from *S.aureus and E.coli* and the quaternary structures of NM23-H1 and *S.aureus* hexameric complexes (Figure 1B). Together, these analyses reveal remarkable conservation of human, prokaryotic and yeast pathogen NDPK proteins across millennia of evolution.

**Figure 1:**
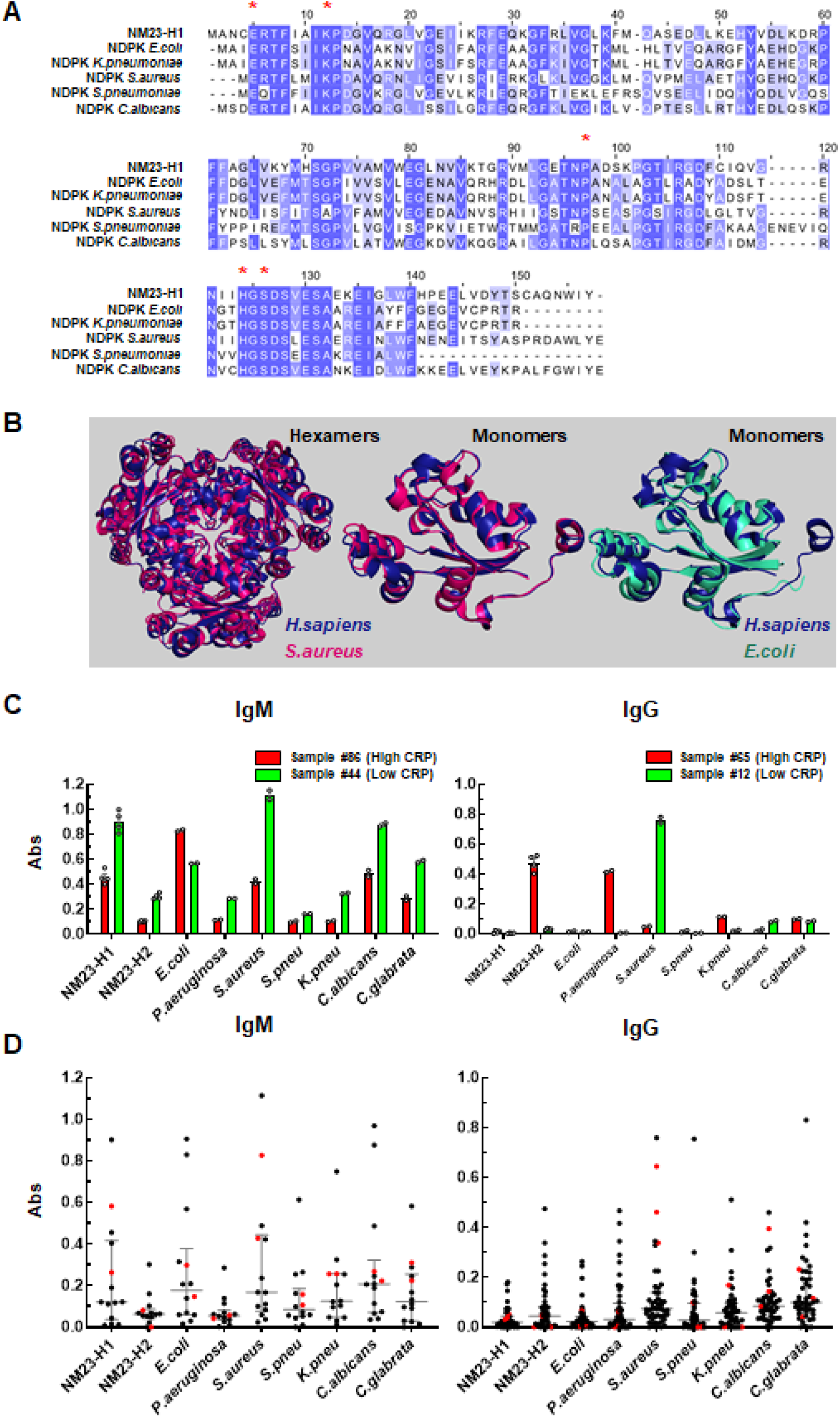
NM23-H1 and pathogen NDPKs are highly conserved and exert the same pro-survival effect on primary AML cells. **A** NDPK protein sequences alignments were obtained with ClustalW and visualised with JalView by Percentage Identity. Red * indicates the key residues for NM23-H1 enzymatic activities and structural organization. **B** 3D alignment of NM23-H1 (blue) crystal structure (pdb 1JXV) to *S.aureus* NDPK (red) tetramer and monomer (pdb 3Q83) and *E.coli* (green) monomer (pdb 2HUR) were generated with PyMol 2.3 (See also Table S1). (C) Representative data of four independent samples (#86 and 44 for IgM; #65 and 12 for IgG), two with high CRP (red bars) and two with low CRP (green bars). CRP values and other samples are summarized in Table S2. Averages are expressed as mean ± SEM. (D) Dot plots for all the analyzed samples; black dots: AML samples; red dots: normal donor samples. Grey bar indicates median ± interquartiles.

### Detection of humoral immunity to pathogen NDPKs

The extensive conservation between human and pathogen NDPKs questions whether the human innate or adaptive immune system can detect and respond to them. However, evidence of humoral immunity to pathogen NDPKs would indicate that exposure and response to pathogen NDPKs occurs *in vivo*. We therefore screened sera from AML patients and normal donors for IgM- and IgG-reactivity against recombinant NDPKs (rNDPKs) from *E.coli, Pseudomonas aeroginosa (P.aueruginosa), S.aureus, S.pneumoniae, K.pneumoniae, C.albicans* and *Candida glabrata (C.glabrata)* and the human NDPKS (NM23-H1 & -H2) (Figure 1C&D). IgM reactivities were tested in 14 AML patients and 2 normal donors. Most donors exhibited IgM activity against a broad spectrum of pathogen and human NDPKs (Figure 1C& D) indicative of B1 B-cell immunity which is fetal in origin and provides natural broad affinity protective IgM anti-PAMP antibodies in the steady state in the absence of antigenic stimulation (35). In contrast, we observed greater variation and selectivity between individuals in IgG reactivity to pathogen NDPKs. For example AML sample #12 displayed selective IgG reactivity with *S.aureus* NDPK, whereas AML sample #65 displayed selective reactivity with *P. aueruginosa* NDPK (Figure 1C). Despite this variation, across a total of 45 AML (selected as described in Supplementary Table S2) and 3 normal donor samples, we identified marked IgG responses to all pathogen NDPKs tested (Figure 1D). NM23-H2 is largely intracellular and serum levels are very low outside of disease. Therefore, these observations suggest that humans experience exposure to pathogen NDPKs *in vivo* and, in doing so, raise antibody repertoires capable of cross-reactivity with human NDPKs which are primarily intracellular during health.

### Pathogen NDPKs recapitulate the actions of NM23-H1 against primary AML cells

We investigated the ability of pathogen NDPKs to recapitulate the actions of NM23-H1 in promoting *in vitro* AML blast cell survival in a significant subset of AMLs (23, 24). We treated 16 individual primary AMLs with rNM23-H1 and pathogen rNDPKs. As shown in Table 1, samples that responded to rNM23-H1 also responded to pathogen rNDPKs, whereas a lack of NM23-H1 response associated with a lack of response to pathogen NDPKs. Seven of 16 (44%) AMLs demonstrated a mean blast cell survival index (BSI; defined as ratio of blasts in NDPK treated cultures versus untreated controls) of >1.5 compared to controls when exposed to rNM23-H1 or pathogen rNDPKs (for absolute cell numbers see Supplementary Table S3). The mean BSI in this group was 4.27 whereas in the non-responders it was 1.03 (p= 0.0047 unpaired t-test with Welch’s correction / 0.0002 Mann Whitney U-test). This response rate is consistent with response rates to NM23-H1 previously published by ourselves and others (23, 24).

**Table 1:**
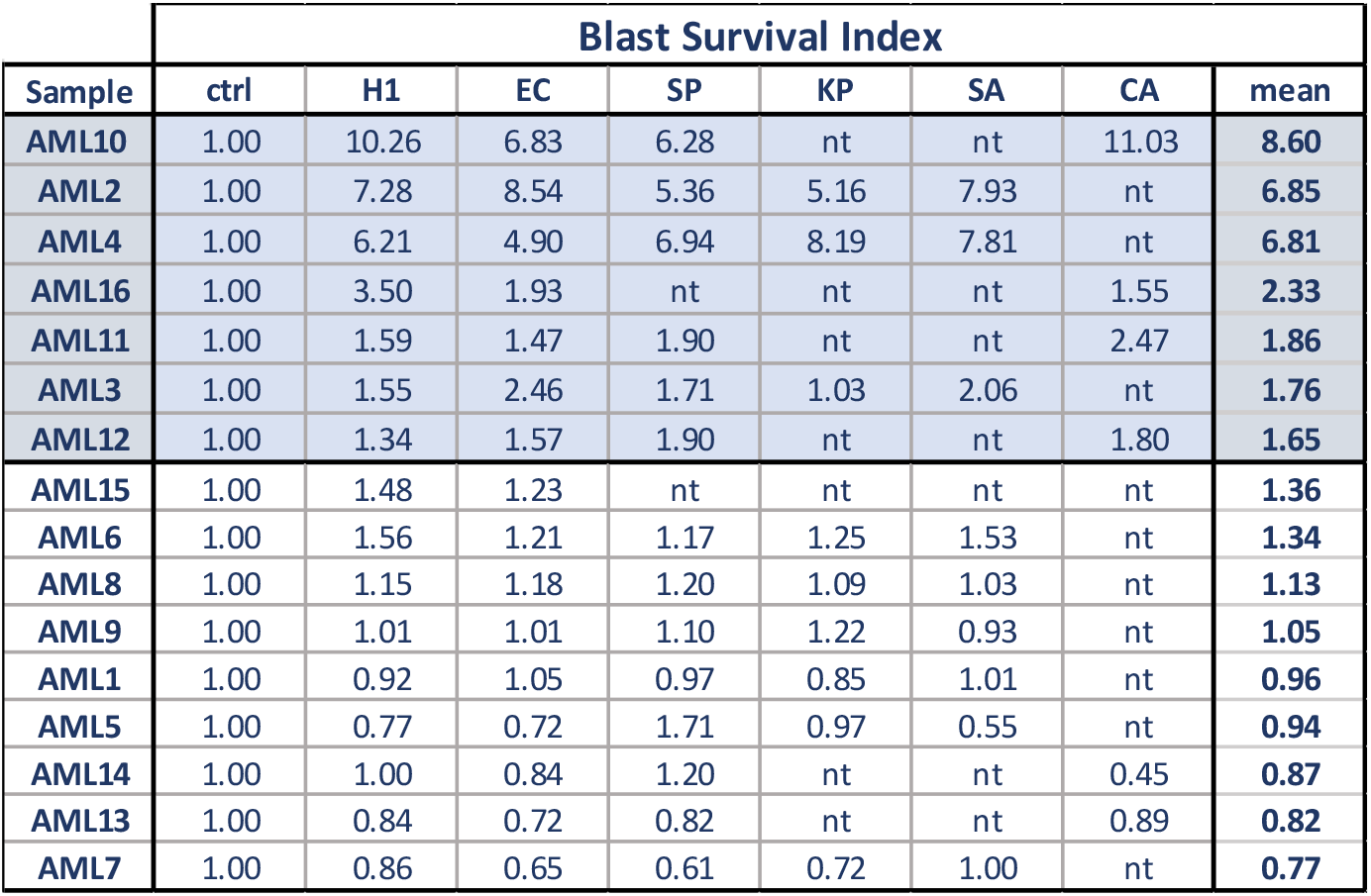
NM23-H1 and pathogen NDPKs are highly conserved and exert the same pro-survival effect on primary AML cells. Primary AML blasts survival index after treatment with rNDPKs for 7 days was calculated relative to vehicle control; mean represent the average response between rNDPKs. Samples were ranked by the mean blast survival index (BSI) and divided into two groups as “responders” (blue background) and “non-responders” (white background) with a cut-off point at BSI > 1.5. H1: rNM23-H1, EC: rNDPK *E.coli*; SP: rNDPK *S.pneumoniae*; KP: rNDPK *K.pneumoniae*; SA: rNDPK *S.aureus*; CA: rNDPK *C.albicans*. nt= not tested

Figure 2 shows in more detail the responses of three selected AMLs (AML1, -2, -4) that displayed the range of responses to rNM23-H1 and pathogen rNDPKs. AML1 demonstrated rNDPK-independent cell survival. This included survival of both AML blasts, identified as CD34^+^ and/or CD117^+^ cells, and of the remaining more mature CD34^-^/CD117^-^ cells in the sample (Figure 2A&B) (See Figure S1 for flow cytometry gating strategy). In contrast, untreated AML2 cells survived poorly *in vitro*, but addition of rNM23-H1 and pathogen rNDPKs equally and markedly increased survival of blasts and more mature cells in the sample (Figure 2A&B). Survival of AML4 in the absence of rNM23-H1 or pathogen rNDPKs was similar to AML1 (Figure 2A&B). However, unlike in AML1, both rNM23-H1 and pathogen rNDPKs equally further increased survival of AML4 blast and mature cells (Figure 2A&2B). In this sample, both rNM23-H1 and pathogen rNDPKs also supported some AML cell proliferation as measured by cells in S+G_2_M phase in flow cytometric cell cycle analyses (Figure 2C). Combined, the data in Table 1 and Figure 2 demonstrate that AML cell responses to pathogen NDPKs are indistinguishable from the previously reported responses to NM23-H1 (23, 24).

**Figure 2:**
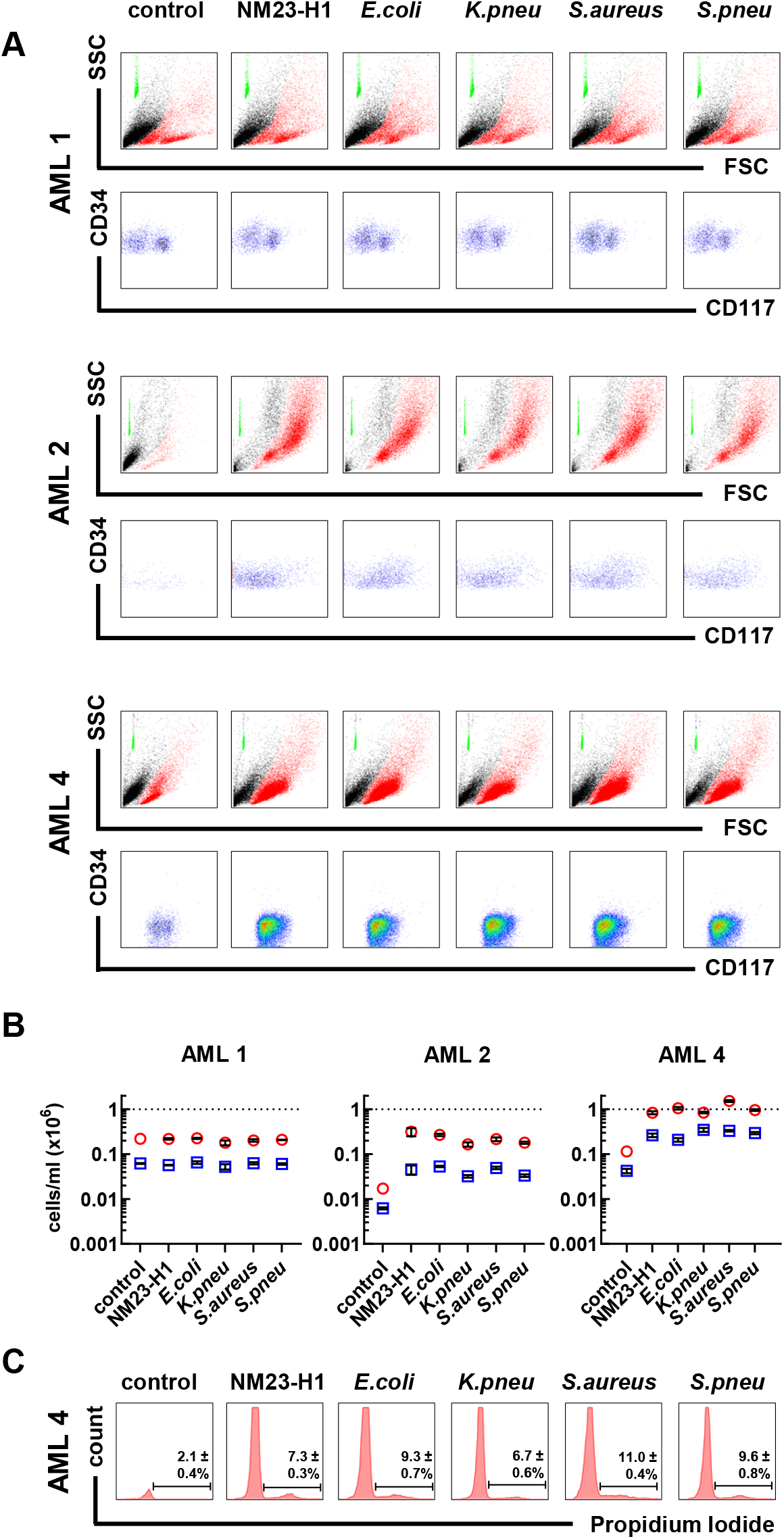
Pathogen NDPKs mimic the action of NM23-H1 against primary AML cells. **A** Representative flow cytometer plots for three primary AML cells treated with rNDPKs for 7 days. Live (in red) and dead cells (in black) were identified by Forward Scatter (FSC) and Side Scatter (SSC). Counting beads (in green) were used to enumerate live cells. Immunostaining with CD34 and CD117 identified the AML blasts amongst live cells. **B** Dot plots show the mean ± SEM of at least 3 replicates for live cells/ml (red circles) and live blasts/ml (blue squares). Dotted line is number of total viable cells at day 0. **C** Flow cytometry histograms of AML 4 cells stained with propidium iodide for cell cycle analysis. Bars and percentages represent cells in S/G_2_M phase. AML 2 and AML 4 treatments are all statistically different to the controls; p<0.05, One-way ANOVA with Dunnet’s test. Gating strategy is presented in supplementary Figure S1

### Promotion of AML cell survival by both human and pathogen rNDPKs is mediated via interaction with non-malignant myeloid cells

We used fluorescently labelled rNM23-H1 and *S.pneumoniae* rNDPK to test for binding to normal donor peripheral blood cells. Both rNM23-H1 and *S.pneumoniae* rNDPK proteins bound to monocytes but not B-cells, T-cells or neutrophils (Figure 3A&B). Notably binding of *S.pneumoniae* NDPK to monocytes appeared less strong than NM23-H1 (Figure 3B, note log scale). We next measured IL-1β and IL-6 production by normal donor peripheral blood cells when exposed to rNM23-H1, *S.pneumoniae* rNDPK and *C.albicans* rNDPK. All three NDPKs induced IL-1β and IL-6 production in PBMC cultures (Figure 3C). However, production of both cytokines was enhanced in cultures of total white cells (TWCs) (Figure 3C) indicating that, although the initial response requires binding of NDPKs to monocytes, the presence of neutrophils in TWC acts to amplify the cytokine response. Whereas at diagnosis AML patients are severely neutropenic, the role of neutrophils in infections during remission might be important. Furthermore, many MDS patients with high risks of AML progression have reasonable numbers of neutrophils that could be involved in pathogen NDPK responses. To further interrogate the importance of monocytes, we exposed leukocytes after red cell lysis to NM23-H1, *E.coli* and *C.albicans* rNDPKs before and after monocyte depletion and measured the resultant secretion of IL-1β and IL-6. As shown in Figure 3D, depletion of monocytes resulted in markedly reduced IL-1β secretion and near abrogated IL-6 secretion. Thus, monocytes are essential and there is interplay between monocytes and neutrophils in the cytokine response to NDPKs.

**Figure 3:**
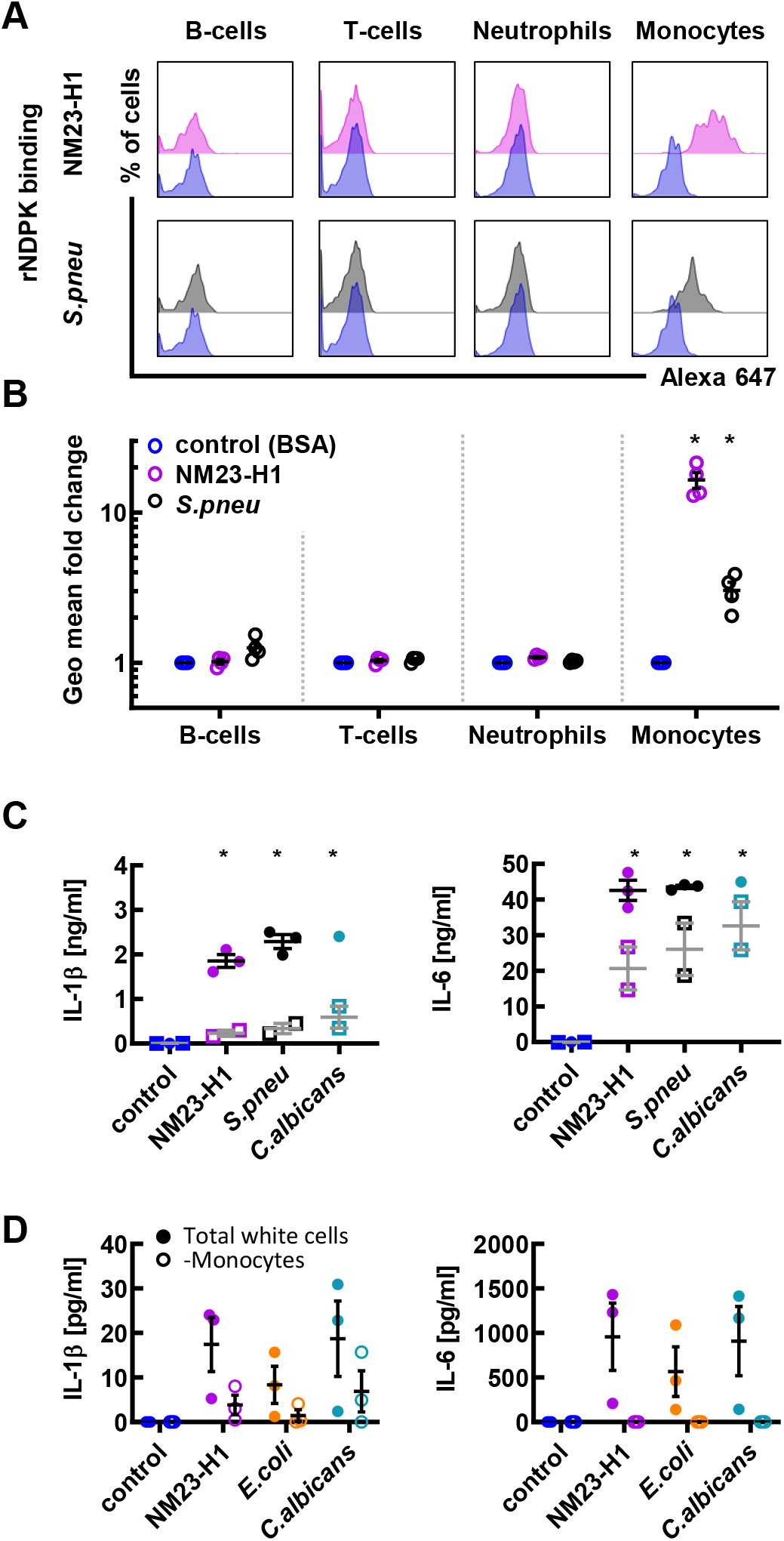
Promotion of AML cell survival by both human and pathogen rNDPKs is mediated via interaction with non-malignant myeloid cells. **A** Whole blood from normal donors was diluted and incubated with fluorescently labelled BSA control (in blue), rNM23-H1 (in pink) and *S.pneumoniae* rNDPKR (in grey). Representative plots for B-cells (CD19), T-cells (CD3), neutrophils (CD11b^+^CD14^-^) and monocytes (CD11b^+^CD14^+^) are shown. **B** Fluorescence geometric mean fold changes compared to BSA control of each individual cell type for n=4 normal donors. **C** ELISA for IL-1β and IL-6 cytokines in conditioned media (CM) generated incubating for 18 hours diluted whole blood (closed circles) or PBMC (open squares) with rNM23-H1 or rNDPK from *S.pneumoniae* and *C.albicans*. **D** Red cells from normal donor’s whole blood were lysed and leukocytes were depleted from CD14^+^ cells (monocytes). Either total white cells or monocyte depleted cells were treated with NM23-H1 or rNDPK for 18h and IL-1β and IL-6 were analyzed by ELISA.

### Neither cytokine induction nor preservation of AML cell viability require enzymatic activity of human or pathogen NDPKs

We used site-directed mutagenesis to make a panel of NTP/NDP transphosphorylase and/or protein histidine kinase deficient rNM23-H1 and *E.coli, S.pneumoniae, C.albicans* rNDPKs and also mutants not able to form higher quaternary structures (tetramers or hexamers) favored by wt NDPKs (summarised in supplementary Table S4). Verification of these mutants is shown in Supplementary Figure S2. Despite having variable enzymatic and oligomerization activities, all the mutant proteins retained the ability to elicit IL-1β secretion from Vitamin D_3_ (VitD_3_) differentiated THP-1 monocytes (Figure 4A) and the ability to enhance primary AML cell survival (Figure 4B).

**Figure 4:**
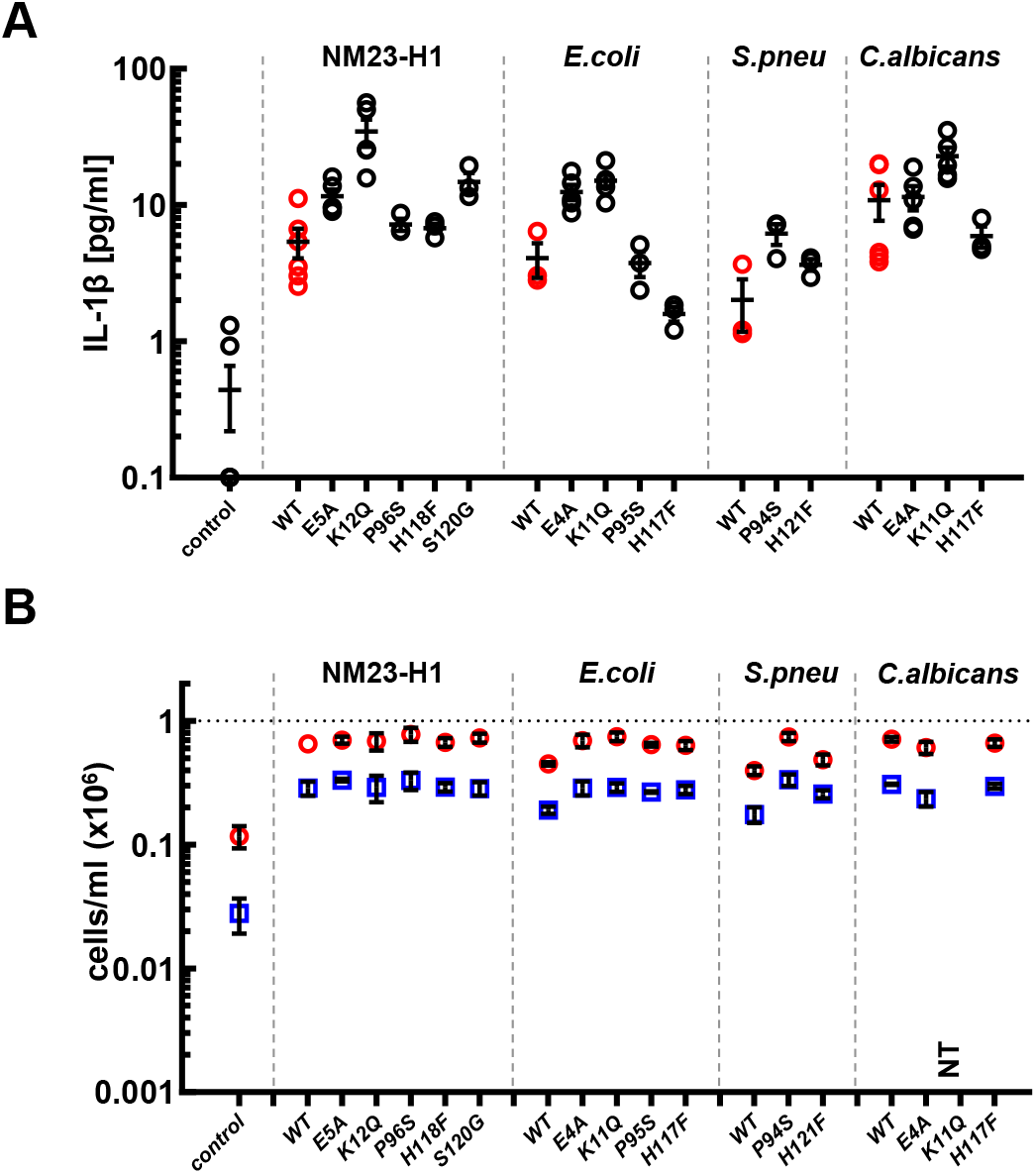
Induction of cytokines and preservation of AML cell viability does not require enzymatic activity of human and pathogen NDPKs. **A** Vitamin D_3_ differentiated THP-1 were incubated for 18 hours in presence of WT (in red) and mutated rNDPK (in black), conditioned media was harvested and IL-1β was measured by ELISA. **B** AML10 was treated for 7 days with WT or mutated rNDPK and cell viability of total cells and blasts was measured by flow cytometry. Dot plots show the mean ± SEM of at least 3 replicates for live cells/ml (red circles) and live blasts/ml (blue squares). All the WT and mutants rNDPK treatments are statistically different from the control; p<0.05, One-way ANOVA with Dunnet’s test. Averages are expressed as mean ± SEM.

### NM23-H1 and pathogen NDPKs activate an NLRP3 inflammasome pathway

IL-1β production by monocytes is intimately associated with activation of the NLRP3 inflammasome which cleaves pro-caspase-1 to release active caspase-1 that in turn cleaves pro-IL-1β, to form active and secreted IL-1β. In many cell types, activation of the NLRP3 inflammasome results in a specialized form of programmed cell death termed pyroptosis (33). However, in human monocytes lipopolysaccharide (LPS) activates an alternative NLRP3 inflammasome pathway that results in IL-1β secretion in the absence of pyroptosis (34).

We used flow-cytometry to measure caspase-1 activation in normal donor leucocytes following exposure to either rNM23-H1, *E.coli* rNDPK or *S.pneumoniae* rNDPK, using LPS and nigericin (an inducer of NLRP3-dependant pyroptosis (36)) as positive controls. All treatments not only induced caspase-1 activation in monocytes, but also in neutrophils (Figure 5A) which do not bind rNM23-H1 protein (Figure 3B). These findings further suggest that neutrophils become activated downstream of NDPKs binding to monocytes and provide the rationale for the amplification of the IL-1β response seen in TWCs compared to PBMC preparations (Figure 3C) and its loss on depletion of monocytes (Figure 3D).

**Figure 5:**
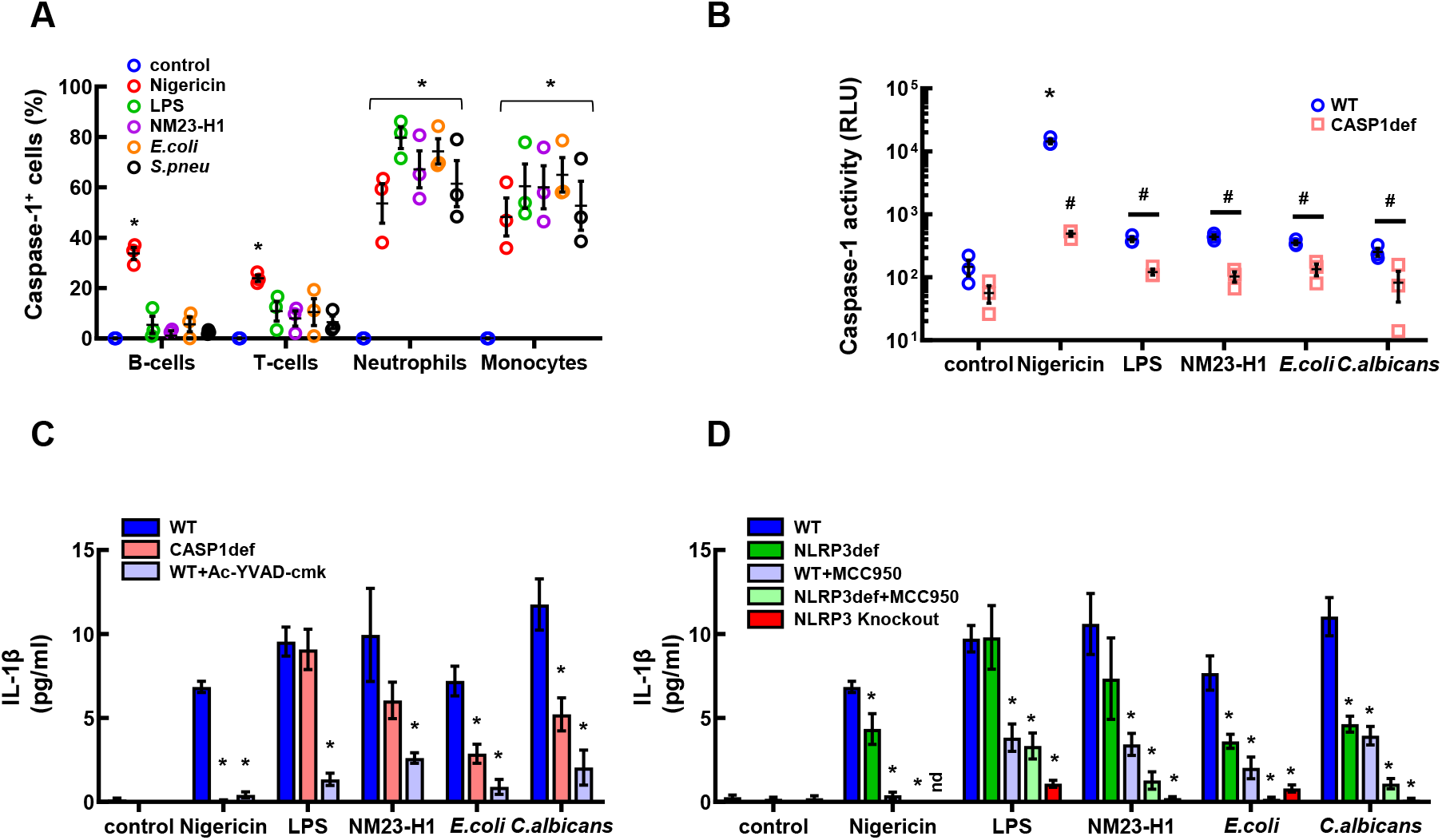
NM23-H1 and pathogen NDPKs activate the NLRP3 inflammasome pathway. **A** Whole blood from normal donors was diluted 2:3 and incubated with nigericin, LPS and rNDPKs. The caspase-1 fluorescent dye FLICA 660 was added after 60’ and incubation continued for further 90’. Red cells were lysed, and leukocytes were immunophenotyped by flow cytometry. * p<0.05 to controls, One-way ANOVA with Dunnet’s test. **B** Caspase-1 specific activity measured with the Caspase-Glo 1 Bioluminescent Inflammasome Assay for both Vitamin D_3_ differentiated THP-1 wild type (WT, blue circles) and Caspase-1 deficient (CASP1def, pink squares) treated for 150’with nigericin, LPS and rNDPKs. Luminescence of the Ac-YVAD-CHO inhibited reactions was subtracted from the total luminescence. **\*** p<0.05 to controls within same cell line, # p<0.05 between WT and CASP1def; One-way ANOVA with post-hoc Tukey’s test. **C** IL-1β ELISA of the conditioned media of Vitamin D_3_ differentiated THP-1 wild type (WT, in blue), in presence or absence of the Caspase-1 inhibitor Ac-YVAD-cmk (light blue), and of the THP-1 Caspase-1 deficient cell line (pink), treated for 18h with nigericin, LPS or rNDPKs. **D** IL-1β ELISA of the conditioned media from Vitamin D_3_ differentiated THP-1 wild type (WT, in blue), THP-1 NLRP3 deficient (NLRP3def, in green) and THP-1 NLRP3 knockout (in red), in presence or absence of the NLRP3 inhibitor MCC950 (light blue and green), treated for 18h with nigericin, LPS or rNDPKs Averages are expressed as mean ± SEM. * p<0.05 to controls, One-way ANOVA with Dunnet’s test.

To further investigate the role of the NLRP3 inflammasome in human monocyte-NDPK responses we used commercially available NLRP3- and caspase-1 deficient (NLRP3def (N_d_), CASP1def (C_d_)), and NLRP3 knockout (N_ko_) THP-1 cell lines and NLRP3- and caspase-1 selective inhibitors. We directly assayed caspase-1 activity in differentiated wtTHP-1 and CASP1def THP-1 cells in the presence or absence of nigericin, LPS, rNM23-H1, *E.coli* rNDPK and *C.albicans* rNDPK (Figure 5B). As expected, baseline caspase-1 activity was lower in CASP1def THP-1 monocytes than in untreated wtTHP-1 monocytes (Figure 5B). Nigericin exposure produced an approximately 1000-fold increase in caspase-1 activity in wtTHP-1 monocytes and this response was significantly reduced in CASP1def THP-1 monocytes (Figure 5B). LPS induced a much smaller increase in caspase-1 activity in wtTHP-1 monocytes and again a reduced response in CASP1def THP-1 monocytes was observed. The induction of caspase-1 activity by human (NM23-H1), bacterial (*E.coli*) and yeast (*C.albicans*) rNDPKs in wtTHP-1- and CASP1def THP-1-monocytes mirrored the response to LPS, and not the response to nigericin (Figure 5B).

The secretion of IL-1β in response to nigericin was abrogated in CASP1def THP-1 monocytes whereas the response to LPS was not affected (Figure 5C). Secretion of IL-1β in response to human (NM23-H1) bacterial (*E.coli*) and yeast (*C.albicans*) rNDPKs showed an intermediate phenotype and was partially diminished in CASP1def THP-1 monocytes compared to wtTHP-1 monocytes (Figure 5C). The caspase-1 selective inhibitor Ac-YVAD-cmk dramatically reduced IL-1β secretion in response to nigericin, LPS and all three rNDPKs (Figure 5C). Collectively, these findings indicate that IL-1β secretion by monocytes in response to NM23-H1 and pathogen rNDPKs is caspase-1 dependent.

VitD_3_-differentiated NLRP3def-THP-1 demonstrated variably reduced IL-1β secretion in response to nigericin, rNM23-H1, *E.coli* rNDPK and *C.albicans* rNDPK whereas the response to LPS appeared unchanged (Figure 5D). Treatment of both wtTHP-1 and NLRP3def-THP-1 monocytes with the NLRP3 inhibitor MCC950 reduced LPS-induced IL-1β secretion (Figure 5D). Similarly, MCC950 markedly diminished IL-1β secretion by wtTHP-1 monocytes in response to nigericin, rNM23-H1, *E.coli* rNDPK and *C.albicans* rNDPK (Figure 5D). In addition, MCC950 further diminished IL-1β secretion by NLRP3def-THP-1 monocytes in response to these stimuli indicating that rNDPK induced IL-1β secretion by monocytes is also NLRP3 dependent. Importantly, IL-1β production, was completely lost in N_ko_ THP-1 cells (Figure 5D).

### NM23-H1 and pathogen NDPK activation of the NLRP3 inflammasome pathway occurs without pyroptosome formation

Active inflammasomes are complexes composed of multiple copies of NLRP3, ASC and pro-caspase-1 proteins. In the classical NLRP3 pathway they ultimately form a single very large aggregate termed the pyroptosome (sometimes referred to ASC specks) which coincide with death by pyroptosis (33). In the more recently described alternative NLRP3-inflammasome pathway, IL-1β secretion by human monocytes occurred in the absence of pyroptosome formation and without pyroptotic cell death (34).

We used VitD_3_-differentiated THP-1 cells stably expressing an NF-κB inducible ASC::GFP fusion gene (ASC::GFP THP-1) to determine whether rNM23-H1 and pathogen rNDPK responses are associated with pyroptosome formation. ASC::GFP expression was induced after 3-6 hours exposure to LPS, rNM23-H1, *E.coli* rNDPK and *S.pnuemoniae* rNDPK (Figure 6A & Figure S3) demonstrating that like LPS, human and pathogen NDPKs activate NF-κB in human monocytes. However, nigericin induces pyroptosis without NF-κB activation. Thus as expected, nigericin treatment alone did not induce ASC::GFP expression (Figure S3). However, pyroptosome formation was observed when nigericin was used to treat LPS exposed (3hr) ASC::GFP expressing cells (Figure 6A & Figure S3). To investigate more closely the induction of pyroptosomes by nigericin in our system, we used ASC-immunofluorescence in wtTHP-1 monocytes. We observed pyroptosome-specks in response to nigericin alone without co-treatment with LPS (Figure 6B&C). However, we again failed to detect increased pyroptosome formation in response to either LPS or rNM23-H1 and rNDPKs (Figure 6B&C). Consistent with the formation of pryroptosomes, nigericin treatment but not LPS, rNM23-H1, *E.coli* rNDPK and *S.pnuemoniae* rNDPK induced rapid cell death of THP-1 monocytes (Figure 6D).

**Figure 6:**
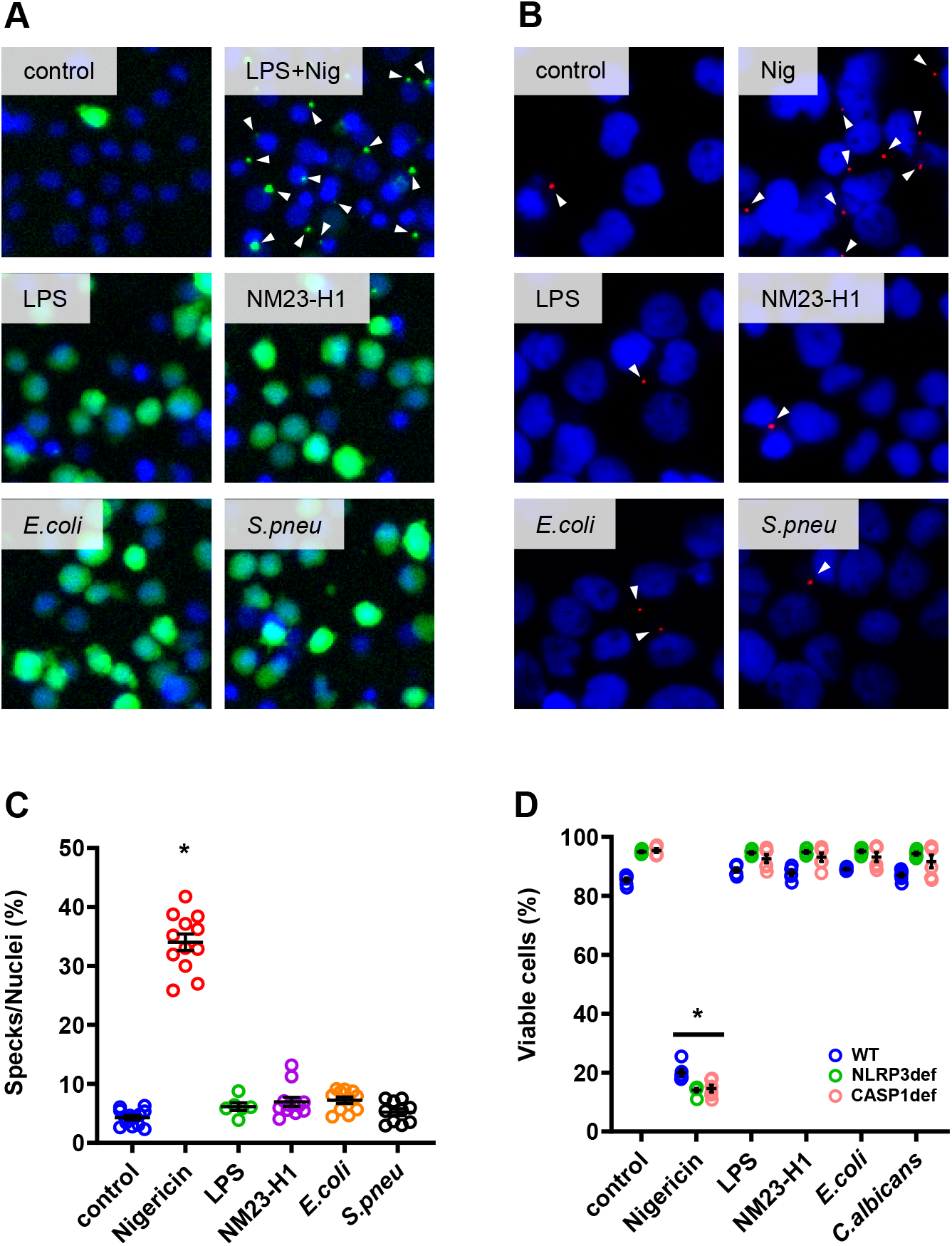
NM23-H1 and pathogen NDPKs activation of the NLRP3 inflammasome pathway occurs without pyroptosome formation. **A** Vitamin D_3_ differentiated THP-1 ASC::GFP cells were treated for 6 hours with LPS and rNDPKs or for 3 hours with LPS to induce ASC::GFP protein with subsequent treatment with Nigericin for further 3 hours (LPS+Nig). ASC::GFP protein was visualized by immunofluorescent microscopy. **B** Immunofluorescent staining of ASC specks in Vitamin D3 differentiated wtTHP-1. Arrows indicate ASC specks. **C** Dot plot of the quantification of the number of ASC specks per total nuclei in THP-1 wild type. * p<0.05 compared to control, One-way ANOVA with Dunnet’s test within the same cell line. **D** Dot plot of the viability of Vitamin D3 differentiated THP-1 WT (WT, in blue), NLRP3 deficient (NLRP3def, in green) and Caspase-1 deficient (CASP1def, in pink) after treatment with nigericin, LPS, and rNDPKs. Viability was measured by flow cytometry and the Forward Scatter/Side Scatter parameters. * p<0.05, One-way ANOVA with Dunnet’s test within the same cell line. Averages are expressed as mean ± SEM.

### Activation of the NLRP3 inflammasome pathway by NM23-H1 and pathogen NDPKs is independent of TLR4 signaling

The production of recombinant proteins in *E.coli* risks contamination with LPS. Although all our experiments using rNDPKs were performed using LPS-neutralizing polymyxin-B (PMB), it was important to rule out the possibility of inflammasome activation by rNDPKS due to low-level LPS contamination. We therefore exploited Dual Reporter THP-1 cells (THP1-Dual) and THP1-Dual TLR4 KO cells (37). These cells have inducible luciferase and SEAP reporter genes, allowing the concomitant study of activation of the IRF and NF-κB pathways, respectively. In addition, THP1-Dual TLR4 KO cells express a truncated TLR4 unable to respond to LPS. For these experiments we also produced human and pathogen rNDPKs in endotoxin-free ClearColi BL 21 cells that contain genetically modified LPS that does not trigger an endotoxic response in humans (38). As in previous experiments we used LPS as a positive control.

As expected, LPS induced NF-κB and IRF activation and secretion of IL-1β in TLR4-competent THP1-Dual monocytes. Responses which were markedly diminished or abrogated by PMB and the TLR4-inhibitor TAK-242 (resatorvid) (39). Also as expected, LPS did not evoke NF-κB and IRF activation or secretion of IL-1β in THP1-Dual TLR4 KO monocytes. In contrast, ClearColi derived rNM23-H1 and pathogen rNDPKs induced NF-κB, IRF and IL-1β responses in both THP1-Dual and THP1-Dual TLR4 KO monocytes when even in the presence of PMB or TAK-242 (Figure 7 A-C) ruling out that these actions were mediated by contaminating LPS. The activation of NF-κB reporter gene expression by NDPKs in THP1-Dual and THP1-Dual TLR4 KO monocytes is consistent with NDPK induction of the NF-κB inducible ASC::GFP fusion gene in ASC::GFP THP-1 monocytes (Figure 6A & Figure S5). Whereas induction of the NF-κB reporter was consistent across the NDPKS tested (Figure 7A), the response of the IRF reporter and secretion of IL-1β were more variable amongst the tested NDPKs (Figure 7 B&C) and correlated with each other (Figure 7 D).

**Fig. 7:**
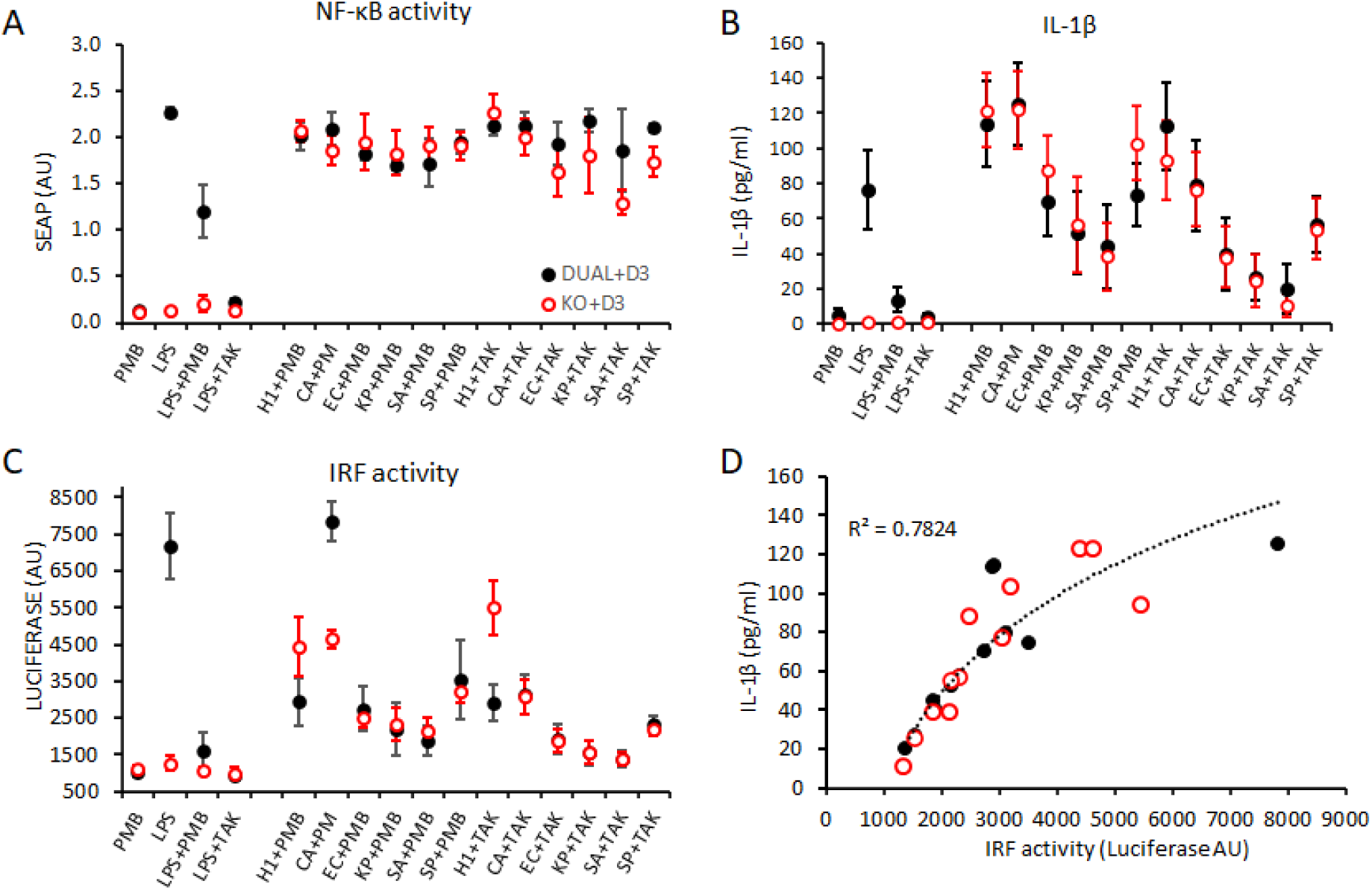
NDPK induced Activation of NF-κB, secretion of IL-1β and IRF activation occur independently of TLR4 signaling. THP1-Dual (black closed circles) and THP1-Dual TLR4 KO (red open circles) were differentiated into monocytes with Vitamin D3. Following 18 hours exposure to the treatments shown, the following were measured: **A** NF-κB-SEAP reporter gene activity. **B** IL-1β ELISA of the conditioned media, and **C** IRF-Lucia-Luciferase reporter gene activity. **D** Data for IRF Lucia-Luciferase reporter gene activity were plotted against IL-1β ELISA results for Vitamin D3-differentiated THP1-Dual (black) and THP1-Dual TLR4 KO (red) monocytes. Line of fit is log^10^. PMB; polymyxin-B. TAK; TAK-242. Data are the mean of n = 4-6 ± SEM.

## Discussion

The demonstration here that human, bacterial and fungal derived NDPKs converge on signaling via a common TLR4-independent NLRP3 inflammasome pathway, has implications for the role of infections in AML and MDS progression. Although a novel concept in these diseases, the association between microbiota and solid cancers is becoming accepted (40-44) and the association of inflammation with cancer is well established (43, 45). In addition, the association of sterile NLRP3 inflammasome activation with AML and MDS is also well accepted (10-16). To date, the role of NLRP3 inflammasomes in myeloid malignancies has focused on classical pyroptotic pathways, where cells dying by pyroptosis release DAMPS that further activate inflammation via TLRs including TLR4 (13, 15, 17, 46-48). However, our study has highlighted the potential of pathogen derived NDPKs acting as PAMPS to activate NLRP3 independently of either TLR4 or pyroptosis. Therefore, our findings widen the potential impact of NLRP3 signaling processes in these settings.

Of note is that many studies of PAMP-associated inflammation utilise LPS as model. However, LPS is restricted to Gram-negative bacterial infections. Here we show that NDPKs from Gram-positive (*S.aureus, S.pnuemoniae*) and Gram-negative (*E.coli, K.pneumoniae*) bacteria and at least one fungus (*C.albicans*), activate inflammatory mechanisms and promote AML cell survival. Furthermore, we demonstrate that although LPS and NDPKs mediate inflammation via NLRP3 and caspase 1 activation, the perception of these distinct classes of PAMP by monocytes is different with LPS being detected by TLR4 and NDPKs not.

A notable finding was the correlation between variable NDPK-induced IL-1β secretion and IRF reporter gene activation in both THP1-Dual and THP1-Dual TLR4 KO monocytes. It is not possible from our data to determine the relationship between IRF signaling and monocyte IL-1β secretion. NM23-H1 was amongst the strongest stimulators of IL-1β secretion and IRF activation whereas *S.pneumoniae* NDPK elicited lower responses. This may reflect differential binding characteristics of different NDKs to human monocytes since we also observed greater binding (∼10 fold) of fluorescently labelled NM23-H1 than *S.pneumoniae* NDPK to healthy donor monocytes.

Although our studies implicate pathogen NDPKs in inflammasome activation, additional studies are required. One question that remains is how monocytes perceive the presence of NDPKs. Future studies should focus on what might be the receptor for NDPKs. Furthermore, although our serological analyses support the argument that humans are exposed to pathogen NDPKs *in vivo*, their role for disease progression *in vivo* and how this is impacted by the presence B1 IgM antibodies now requires investigation.

## Materials and Methods

### Ethics

Primary samples were obtained under ethical approval: University of Birmingham, ERN_17-0065 for healthy blood samples and by NRES Committee North West - Cheshire REC reference 12/NW/0742 for AML bone marrow aspirates.

### Cell lines and tissue culture

THP-1 (DSMZ) and THP-1 NLRP3 deficient (N_def_), THP-1 Caspase-1 deficient (Casp1_def_), THP-1 NLRP3 knockout (N_ko_), THP-1 ASC::GFP, THP1-Dual, and THP1-Dual-TLR4 KO Cells (all Invivogen) were maintained between 0.5-1×10^6^ cells/ml in RPMI 1640 media, supplemented with 10% Fetal Bovine Serum and 100U/ml penicillin, 100μg/ml streptomycin (pen/strep) (complete media) at 37°C, 5% CO_2_ in a humidified incubator. To select for reporter gene expression THP1-Dual and THP1-Dual TLR4 KO cultures were also supplemented with 100µg/ml Zeocin and 10µg/ml Balsticidin. Cells were STR-profiled regularly throughout the study.

### DR THP1 and TLR4 KO DR THP-1 Cell reporter assays

THP1-Dual™ and TLR4 KO THP1-Dual™ cells feature the secreted Lucia luciferase reporter gene, under the control of an ISG54 (interferon-stimulated gene) minimal promoter in conjunction with five interferon (IFN)-stimulated response elements. The same cells also express a secreted embryonic alkaline phosphatase (SEAP) reporter gene driven by an IFN-β minimal promoter fused to five copies of the NF-κB consensus transcriptional response element and three copies of the c-Rel binding site. These reporter proteins were measured in cell culture supernatants using QUANTI-Luc™ (IRF activity) and QUANTI-Blue™ (NF-κB activity) assays as described by the manufacturer (Invivogen).

### Protein production

The complete coding sequences of NM23-H1, bacterial and fungal NDPKs were cloned into a pET15b backbone (Merck Millipore) and transformed into BL21 (DE3) or ClearColi BL21 (DE3) bacteria. Protein expression was induced with 1mM Isopropyl β-D-1-thiogalactopyranoside (IPTG) overnight at 20°C in a shaking incubator. Recombinant proteins were purified with His-Bind Resin and His-Bind Buffers (Merck Millipore) according to manufacturer’s protocol and stored at a concentration of 2mg/ml in elution buffer (Merck Millipore) at 4°C or diluted to 0.2mg/ml in culture media and filter sterilized for cell treatments.

### Indirect ELISA for anti-NDPK IgG and IgM

Nunc MaxiSorp flat-bottom ELISA Plates (ThermoFisher Scientific) were coated with 100µl of rNDPK at 1µg/ml in PBS overnight at 4°C or PBS alone. Plates were then washed for 4 times with 300µl of PBS-Tween-20 (0.2% Tween-20, PBS-T) and blocked with 200µl of 2% BSA in PBS for 1 hour at room temperature, shaking at 500rpm. Plates were then washed once with PBS-T and incubated with 100µl of patient plasma/serum diluted 1:200 in PBS-T for 1 hour at room temperature whilst shaking. Plates were washed 4 times with PBS-T and incubated with 100µl of HRP-labelled specific α-human IgG (Clone R10) and α-human IgM (Clone AF6) antibodies diluted 1:1000 in PBS for 1 hour at room temperature whilst shaking. Plates were washed 5 times with PBS-T and 100µl of 3,3′,5,5′-Tetramethylbenzidine (TMB) (Merck Millipore) equilibrated at room temperature were added, and reaction was blocked after 15 minutes with 2N H_2_SO_4_. Plates were read at 450nm with VICTOR X3 Light Plate Reader (PerkinElmer). Absorbance was then calculated by subtracting the cross-reactivity of HRP-labelled detection antibodies against rNDPK and baseline obtained for blocking solution.

### Primary AML direct survival assay

Bone marrow leukocytes were washed in RPMI 1640 supplemented with pen/strep (Gibco), and resuspended in RPMI 1640 supplemented with 1% v/v ITS+ Premix (Corning) and pen/strep (Serum-free media) at 1×10^6^ cells/ml. Treatments were performed with 2μg/ml of rNM23-H1/rNDPKs, in the presence of 1.25μg/ml polymyxin B (Merck Millipore) (see also supplementary Figure S2). Cells were incubated for 7 days and immunostained for flow cytometry (Antibodies: CD34, clone 581; CD117, clone: 104D2; CD11b, clone ICRF44 (BD Pharmingen) and 1:50 FcR Blocking Reagent (Miltenyi Biotec)). Cells were analyzed on a FACS Calibur (BD Pharmingen) in the presence of counting beads for live cells enumeration.

### Cell cycle analysis

Cell cycle was assessed by incubating 300µl of cells with 200µl of Cell Cycle Buffer (30μg/mL Propidium Iodide (PI), 0.1mM NaCl; 1% Triton X-100) for 24 hours at 4°C in the dark and before analysis on a BD FACS Calibur.

### rNDPK Alexa 647 labelling

Purified rNM23-H1, rNDPK form *S.pneumoniae* and Bovine serum albumin (BSA) were diluted to 2mg/ml and dialyzed at 4°C overnight in 2000 volumes of 0.5M NaCl, 0.1M imidazole in phosphate buffered saline (PBS), pH 8.2, prior to labelling using the Alexa Fluor 647 Protein Labelling Kit (ThermoFisher Scientific) according to manufacturer’s instructions.

### rNDPK Binding to normal leukocytes

Normal adult donor blood (in EDTA) was diluted 1:5 with RPMI 1640 supplemented with pen/strep and 2mM L-glutamine (Gibco). Cells were incubated at 37°C for 2 hours in the presence of polymyxin B (1.25μg/ml) and either 2μg/ml of Alexa 647 labelled NDPKs or BSA. Red cells were then lysed with red cell lysis buffer (155mM ammonium chloride, 10mM potassium bicarbonate, 0.1mM EDTA, pH 8.0) and leukocytes immunostained for CD19 (clone SJ25C1) CD3 (clone SK7), CD11b (clone D12), and CD14 (clone M5E2) (all BD Pharmingen), in the presence of 1:50 FcR Block (Miltenyi Biotec) before flow cytometric analysis.

### Differentiation of THP-1 monocyte models

Prior to treatments, THP-1 cell lines were diluted to 0.2×10^6^ cells/ml and differentiated with 100nM 1α,25-dihydroxy Vitamin D_3_ (Cayman Chemical) for 72 hours. Differentiation was confirmed by measuring expression of CD11b and CD14 markers by flow cytometry (antibodies: CD11b, clone D12; CD14, clone M5E2, BD Pharmingen).

### Normal donor mononuclear preparation

Peripheral Blood Mononuclear Cells (PBMC) from normal adult donor blood were isolated using Ficoll-Paque PLUS (Fisher Scientific). After purification, cells were resuspended at 1×10^6^ cells/ml in 1% ITS+ RPMI 1640 media for the treatments.

### Monocyte depletion

Red cells from normal adult donor blood (in EDTA) were lysed in red cell lysis buffer and leukocytes were rested for 24 hours in a non-coated flask in RPMI 1640 media supplemented 10% autologous plasma. After 24 hours, total leukocytes were depleted from monocytes with CD14 MicroBeads UltraPure (Miltenyi Biotec) according to manufacturer’s instruction and treated in autologous media.

### Cytokine analysis

Diluted whole blood (1:5), 1×10^6^ PBMC/ml, total leukocytes, monocyte-depleted leucocytes, or Vitamin D_3_ differentiated THP-1/ml were treated with 2μg/ml of rNM23-H1 or rNDPKs, and when possible with 2μg/ml of ultrapure lipopolysaccharide from *E.coli* K12 (Invivogen) and 20μM Nigericin (Merck Millipore) as control for alternative and canonical inflammasome, respectively. Polymyxin B (PMB) 1.25μg/ml was used for all the treatments except LPS. Cells were incubated for 18 hours and conditioned media (CM) collected by centrifugation and stored at -20°C. IL-1β and IL-6 were analyzed by ELISA MAX Deluxe Sets (Biolegend) according to manufacturer’s instruction.

### Caspase-1 activation

#### Normal leukocytes

Normal adult donor blood (EDTA) was diluted 2 + 3 in RPMI160. Cells were treated as for the cytokine analysis for one hour with Nigericin, LPS, rNM23-H1 and rNDPK and incubated at 37°C. FLICA 660 Caspase-1 (Bio-Rad) substrate was then resuspended in DMSO following the manufacturer’s protocol and applied 1:30 on the cells. Non-FLICA treated cells were vehicle controlled. Cells were then incubated for a further 1.5 hours with gentle pipetting every 30 minutes. Red cells were then lysed and leukocytes immunophenotyped by flow cytometry analysis as described above. Caspase-1 positive cells were calculated with Population Comparison (SE Dymax) in FlowJo 10.6. Version 10.6.0.

#### THP-1 cell lines

Cells were plated and treated as for the cytokine analysis, but incubated for 2.5 hours. CM was then harvested by centrifugation and active caspase-1 measured with the Caspase-Glo 1 Inflammasome Assay (Promega) following the manufacturer’s instructions. Luminescence was measured after 90’ with VICTOR X3 Light Plate Reader (PerkinElmer). Luminescence values obtained in the presence of Ac-YVAD-cho inhibitor were subtracted from the non-inhibited reaction, to obtain the caspase-1-specific activation.

### Inhibition of inflammasome components

Inhibitors and antagonists used were: Caspase-1 inhibitor Ac-YVAD-cmk (Invivogen), 18.5μM in DMSO; NLRP3 inhibitor MCC950, 10μM in DMSO; TLR4 inhibitor TAK242 2.76µM in DMSO. Cells were pre-treated for 1 hour with the inhibitors, while TLR4 antagonist was added from the beginning of the incubation period.

### ASC Speck/Pyroptosome formation

Pyroptosome formation (ASC speck) was investigated both with THP-1 ASC::GFP cell line (Invivogen) and THP-1 wild type. Cells were differentiated with Vitamin D_3_ prior to treatment. THP-1 ASC::GFP nuclei were stained with 250ng/ml Hoescht 33342 (Merck Millipore) for 45 minutes, washed and resuspended at 0.5×10^6^ cells/ml in phenol red free complete media and treated with 2μg/ml of LPS or rNDPK. For nigericin treatment, since it cannot induce the NfκB-dependent fusion gene ASC::GFP, it was necessary to pre-treat cells with 2μg/ml LPS and then with 20μM Nigericin (LPS+Nig) in order to induce canonical inflammasome and visualize pyroptosomes (ASC specks). THP-1 wild type were used to immunostain for ASC specks. Cells were treated for 6 hours and cytospun on glass slides. After fixation with 4% paraformaldehyde, cytospins were blocked with 10% heat inactivated goat serum (Merck Millipore), 1% FBS (Gibco) in PBS (Gibco) for 30 minutes at 37°C and then incubated with 10 µg/ml of αASC TMS-1 (clone HASC-71, Biolegend) in blocking buffer, overnight, at 4°C. Texas Red labelled secondary antibody (Jackson ImmunoResearch) was used at 1:200, for 1 hour, at 37°C. Micrographs were acquired with EVOS FL (ThermoFisher Scientific) and analysis was performed with the ImageJ distribution (49).

### Primary AML indirect survival assay

CM was prepared from normal donor 1:5 diluted whole blood or 1×10^6^ PBMC/ml +/- rNDPK and rNDPK depleted as described before (24). AML cells from 3 primary AML bone marrow aspirates were resuspended in CM at 1×10^6^ cells/ml, incubated for 7 days and viability assessed with flow cytometry and counting beads.

### Quantification and Statistical Analysis

Results are expressed as means ± standard error (SEM) from at least three independent replicates for each experimental group. Statistical analyses were performed with GraphPad Prism 9 and the relevant test used is specified in the Figure legends. Values of p < 0.05 are identified with an *.

## Acknowledgements

This research was supported by a grant from Blood Cancer UK (Grant 17011). SL has been supported by the state of Baden-Württemberg within the PharmCompNet project. We are grateful to Dr Adrian Shields for critical review of the manuscript.

## List of abbreviation

AML: acute myeloid leukemia
ASC: apoptosis-associated speck-like protein containing a caspase-recruitment domain
CEBPa: CCAAT/enhancer-binding protein alpha
COD: cause of death
DLBCL: diffuse large B-cell lymphoma
IL: interleukin
IL-1Ra: interleukin-1 receptor antagonist
LPS: lipopolysaccharide
MDS: myelodysplastic syndrome
NDPK: nucleoside diphosphate kinase
NLRP3: NOD-, LRR- and pyrin domain-containing protein 3
NTP/NDP: nucleoside tri/diphosphate
PAMP: pathogen associated molecular pattern
PRR: pattern recognition receptor
rNDPK: recombinant NDPK
TBC: total blood cells
TLR4: toll-like receptor 4

## Author contributions

CMB, FLK and MTD conceived the hypothesis and supervised the project. ST conceived, planned and performed experiments throughout the paper. FL, MF, BC, AA, AP contributed to AML treatment experiments. BC and AA contributed to evaluation of IL-1β secretion in normal blood and THP-1 models. DS and KD contributed to the experiments using THP1-Dual and TLR4 KO THP1-Dua cells. SL and TW designed and performed auto phosphorylation and kinase activity assays. MG assisted in developing the assays for humoral IgM and IgG. DW DPW selected AML samples for humoral IgM and IgG from archived samples (Manchester Cancer Research Centre Biobank) under the academic supervision of DW and management by DPW. All authors provided critical feedback and/or helped shape the research, analysis and manuscript. CMB, ST and FK, wrote the manuscript.

